# Antagonism between sugar-dependent *O*-fucosylation and BIN2-mediated phosphorylation regulates metabolism and growth in Arabidopsis

**DOI:** 10.64898/2026.11.06.764934

**Authors:** Hongliang Zhang, Yalikunjiang Aizezi, Ajeet Chaudhary, Zhi-Yong Wang

**Affiliations:** Department of Plant Biology, Carnegie Institution for Science, Stanford, CA 94305, United States; Department of Genetics, Stanford University, Stanford, CA 94305, United States; Department of Biology, Stanford University, Stanford, CA 94305, United States

**Author notes:** Corresponding authors: Zhi-Yong Wang and Hongliang Zhang. Department of Plant Biology, Carnegie Institution for Science, Stanford, CA 94305, United States.

## Abstract

Coordinating sugar sensing, hormone signaling, and carbon metabolism is crucial for plant growth and environmental acclimation. Brassinosteroid (BR)-dependent phosphorylation and sugar-dependent *O*-glycosylation modify overlapping sets of proteins, but how these modifications co-regulate protein functions remains unknown. Here, we show that SPINDLY (SPY)-catalyzed *O*-fucosylation competes with BIN2-mediated phosphorylation and antagonistically regulates phosphoenolpyruvate carboxykinase 1 (PCK1), a key gluconeogenic enzyme. SPY associates with and *O*-fucosylates PCK1 within a peptide containing Ser-62 and Thr-66, two BIN2 phosphorylation sites that regulate PCK1 activity, preventing its phosphorylation by BIN2. PCK1 *O-*fucosylation and phosphorylation exclude each other and oppositely affect PCK1 activity *in vivo*. The *spy* mutants show increased PCK1 phosphorylation, reduced PCK activity, and impaired hypocotyl elongation; BR or BIN2 inhibitor treatments suppressed these phenotypes, indicating that these *spy* effects result from increased BIN2 phosphorylation. Together, these results reveal functional antagonism between SPY/*O*-fucosylation and BIN2/phosphorylation in regulating metabolism and growth.

## Introduction

Plant growth requires coordination of developmental signals with the availability and use of carbon. Growth-promoting hormones stimulate cell division, expansion, and differentiation, but these processes can proceed only when sufficient sugars provide energy, metabolic intermediates, and substrates for cell wall biosynthesis (Zhang et al., 2016). Conversely, carbon availability must be translated into appropriate developmental outputs rather than merely increasing metabolic flux. In addition, non-growing cells must maintain an appropriate basal level of carbon metabolism to sustain cellular function and viability. Plants therefore require mechanisms that integrate hormonal information about when and where to grow, or to not grow, with metabolic information and metabolic activities.

Brassinosteroids (BRs) are central regulators of plant growth and act through the receptor kinase BRASSINOSTEROID INSENSITIVE 1 (BRI1) to inhibit BRASSINOSTEROID-INSENSITIVE 2 (BIN2), a glycogen synthase kinase 3 (GSK3)-like kinase. BIN2 is best known for phosphorylating and inhibiting the BRASSINAZOLE RESISTANT 1 (BZR1)/BRI1 EMS SUPPRESSOR 1 (BES1) family transcription factors, thereby coupling BR perception to broad transcriptional programs (He et al., 2002; Li and Nam, 2002; Kim et al., 2009). However, the expanding repertoire of BIN2 substrates indicates that BR signaling also directly regulates diverse cellular processes, including primary metabolism (Kim et al., 2023; Zhang et al., 2026). How hormone-regulated phosphorylation is coordinated with sugar status remains poorly understood.

One major mechanism by which eukaryotic cells sense nutrient availability is the *O*-glycosylation of nucleocytoplasmic proteins (Hart et al., 2007; Lagerlöf et al., 2016; Bi et al., 2023). In Arabidopsis, two related glycosyltransferases catalyze distinct forms of this modification: SECRET AGENT (SEC) transfers *O*-linked N-acetylglucosamine (*O*-GlcNAc) from UDP-GlcNAc, whereas SPINDLY (SPY) transfers *O*-linked fucose (*O*-fucose) from GDP-fucose to serine and threonine residues (Zentella et al., 2016; Zentella et al., 2017; Bi et al., 2023). Because these nucleotide-sugar donors are generated through central carbohydrate metabolism, protein *O*-glycosylation provides a direct route for cellular sugar status to influence protein function (Hart et al., 2007; Aizezi et al., 2025). Genetic studies show that SEC and SPY have overlapping essential functions, as well as distinct roles in hormone signaling, development, circadian regulation, and stress responses (Hartweck et al., 2002; Xu et al., 2017; Bi et al., 2021; Aizezi et al., 2025). SPY is also required for sugar-dependent seedling growth, supporting a physiological role for *O*-fucosylation in coupling carbon availability to growth (Bi et al., 2023). At the molecular level, *O*-glycosylation can alter protein conformation, stability, activity, localization, or interactions, allowing nutrient information to regulate cellular processes rapidly and independently of transcription (Hart et al., 2007; Aizezi et al., 2025).

Modification of serine and threonine residues by either sugars or phosphate creates extensive potential for crosstalk between *O*-glycosylation and phosphorylation (Hart et al., 2011). The two modifications may compete for the same residue, influence modification at nearby residues, or exert opposing effects on protein structure and activity. Antagonistic interactions between *O*-GlcNAcylation and phosphorylation have been observed in animals (Hart et al., 2011), but their functional importance in plants remains largely unexplored (Aizezi et al., 2025). Recent proteomic studies have identified hundreds of *O*-GlcNAcylated and *O*-fucosylated proteins as SEC and SPY substrates (Xu et al., 2017; Bi et al., 2023; Zentella et al., 2023; Shrestha et al., 2024). Independently, proximity labeling combined with quantitative phosphoproteomics identified a broad BIN2 signaling network containing numerous *in vivo* kinase substrates (Kim et al., 2023). Comparison of these datasets revealed substantial overlap between proteins targeted by *O*-glycosylation and BIN2-dependent phosphorylation (Kim et al., 2023; Aizezi et al., 2025). Whether sugar-dependent *O*-glycosylation and BIN2 phosphorylation directly modulate each other on shared substrates, and whether such crosstalk integrates metabolic and BR signals, remain important unanswered questions.

A particularly compelling shared target is PHOSPHOENOLPYRUVATE CARBOXYKINASE 1 (PCK1, also termed PEPCK1) (Xu et al., 2017; Kim et al., 2023), which catalyzes the ATP-dependent decarboxylation and phosphorylation of oxaloacetate to produce phosphoenolpyruvate, a key step in gluconeogenesis. This reaction enables carbon derived from storage lipids, amino acids, and organic acids to enter carbohydrate biosynthesis (Leegood and Walker, 2003; Walker et al., 2021). PCK1 is especially important during post-germinative growth, when seedlings must convert stored reserves into soluble sugars before photosynthesis is fully established. Arabidopsis *pck1* mutants accumulate less soluble sugar and show severe defects in hypocotyl growth and seedling establishment under carbon-limiting conditions (Rylott et al., 2003; Penfield et al., 2004; Zhang et al., 2026). Beyond early seedling development, PCK functions as a major decarboxylase in PCK-type C_4_ photosynthesis and in some crassulacean acid metabolism plants (Burnell and Hatch, 1988; Walker and Leegood, 1996; Leegood and Walker, 2003). PCK regulation therefore provides a strategic point for matching sugar production with developmental and environmental demands.

A recent study showed that BIN2 directly phosphorylates and inhibits PCK1, whereas BR signaling promotes PCK1 dephosphorylation and activation, establishing PCK1 as a direct link between a growth-hormone pathway and carbon metabolism (Zhang et al., 2026). Its identification as an *O*-glycosylated protein suggests that sugar status and BR signaling converge on PCK1 through interacting post-translational modifications (Xu et al., 2017). Here, we investigate whether SPY-mediated *O*-fucosylation modulates BIN2-dependent phosphorylation of PCK1 and how this interplay affects PCK activity and seedling growth.

## Materials and methods

### Plant materials and growth conditions

*Arabidopsis thaliana* Columbia-0 (Col-0) was used as the wild type. The *spy-4 allele* (Swain et al., 2001), *spy-23* (WiscDsLox241C03) (Woody et al., 2007; Bi et al., 2023), *bzr1-1D*, the semidominant gain-of-function mutant *bin2-1*, and the 35S:PCK1-YFP/Col-0 and PCK1-YFP/*bin2-1* lines (Zhang et al., 2026) were described previously. The *35S:PCK1-YFP/spy-4* line was generated by crossing 35S:PCK1-YFP/Col-0 with *spy-4*. A homozygous line was identified in the F2 generation by screening for Basta resistance and the *spy-4* phenotype and propagated to the F3 generation for experiments. Seeds were surface-sterilized with chlorine gas for 2 h and sown on 120-mm square polystyrene Petri dishes (Greiner Bio-One) containing 0.5× Murashige and Skoog (MS) salts (Duchefa) with the indicated concentrations of sucrose and chemical treatments. Plates were stratified at 4 °C in the dark for 3 d and then grown at 22 °C under a 16-h light/8-h dark photoperiod. For etiolated seedling cultivation, plates were exposed to light for 6 h to synchronize germination before incubation at 22 °C in continuous darkness for the indicated periods. *Nicotiana benthamiana* plants were cultivated in a greenhouse at 20 °C under a 16-h light/8-h dark cycle; approximately 6-week-old plants were used for *Agrobacterium*-mediated infiltration.

### Chemical treatments

Half-strength MS medium was supplemented with the indicated concentrations of propiconazole (PPZ), brassinolide (BL), 30 μM bikinin, or 30 μM SOFTI (Aizezi et al., 2024); corresponding solvent controls were included.

### Hypocotyl length measurements

Seedlings were photographed using a digital camera (Nikon D600). Hypocotyl length of seedlings was measured with ImageJ software.

### PCK activity assays

Measurement of PCK activity in crude plant extracts was performed as described previously (Zhang et al., 2026). In brief, harvested plant tissues were ground to fine powder in liquid nitrogen. Ground tissue (20 mg) was extracted with 500 μl of 100 mM Bicine-KOH (pH 9.0) containing 10% (v/v) glycerol, 0.1% (v/v) Triton X-100, 5 mM β-mercaptoethanol, 1 mM EDTA, 1 mM EGTA, 2 mM phenylmethylsulfonyl fluoride (PMSF), 1× protease inhibitor cocktail (Roche, 11873580001), and 1× phosphatase inhibitor (Pierce, A32957). The extracts were vortexed and centrifuged at 14,000 × g for 10 min. PCK decarboxylation activity in the supernatant was determined at 25 °C by monitoring NADH oxidation at 340 nm. The reaction mixture contained 65 mM Tris-acetate (pH 7.4), 100 mM KCl, 0.3 mM oxaloacetate, 1 mM ATP, 10 μM MnCl₂, 4 mM MgCl₂, 71.5 mM β-mercaptoethanol, 0.4 mM NADH, 1 U pyruvate kinase, and 2 U lactate dehydrogenase. One unit of PCK activity was defined as the amount of enzyme catalyzing the oxidation of 1 μmol NADH min⁻¹ at 25 °C.

### Protein extraction and immunoblotting

Protein extraction and immunoblotting were performed as described previously (Zhang et al., 2025). Briefly, plant material was ground in liquid nitrogen and extracted in Laemmli buffer [125 mM Tris-HCl (pH 6.8), 10% (v/v) 2-mercaptoethanol, 0.01% (w/v) bromophenol blue, 4% (w/v) SDS, and 20% (v/v) glycerol] and incubated at 95 °C for 10 min. Protein samples were separated by standard SDS-PAGE or Phos-tag SDS-PAGE and transferred to polyvinylidene difluoride (PVDF) membranes by semi-dry transfer. Membranes were blocked with 5% (w/v) low-fat milk powder (AppliChem) in TBS-T (0.1% Tween-20) for 1 h at room temperature and probed with anti-PCK1 [1:5,000; (Rojas et al., 2021)], anti-GFP (1:2,000; HT801, Transgen Biotech), anti-His (1:3,000; 27E8, Cell Signaling Technology), anti-mCherry/RFP (1:5,000; ab213511, Abcam), or anti-GST (1:3,000; 26H1, Cell Signaling Technology), as indicated. After three 10-min washes in TBS-T, membranes were incubated with horseradish peroxidase-conjugated goat anti-rabbit or goat anti-mouse IgG (1:5,000; Fisher Scientific) and washed three times for 10 min in TBS-T. Signals were detected using WesternBright Quantum (Advansta). For the nuclear protein immunoblotting, nuclear protein fractions from WT, *bin2-1*, *spy-4*, and *spy-23* seedlings were probed with a mixture of anti-serine-*O*-fucose and anti-threonine-*O*-fucose antibodies (1:1,000; GlycoScientific). Coomassie Brilliant Blue staining or the indicated housekeeping protein served as a loading control.

### AAL affinity purification and AAL-biotin detection

AAL-agarose affinity purification was used to enrich fucose-containing proteins from agro-infiltrated *N. benthamiana* leaves and *Arabidopsis* seedlings. Around 250 mg infiltrated *N. benthamiana* leaf tissue or *Arabidopsis* tissue was homogenized in 1.5 mL of extraction buffer containing 50 mM Tris-HCl (pH 7.5), 500 mM NaCl, 0.5% (v/v) Triton X-100, 2.5 mM 2-mercaptoethanol, 20 μM MG132, and 1× cOmplete EDTA-free protease inhibitor cocktail (Sigma-Aldrich). All samples were sonicated for 1 min on ice and centrifuged at 15,000 g for 10 min at 4°C to clarify the extracts. An aliquot of each clarified extract was retained for input analysis. 20 μL of extract was mixed with 20 μL of 2× Laemmli sample buffer, and the remaining extract was incubated with 30 μl of AAL-agarose beads (2 mg lectin/ml, AL-1393-2; Vector Laboratories) for 1.5 h at 4°C with end-over-end rotation. Beads were washed by four washes with TBST-500 and one wash with 20 mM Tris-HCl (pH 7.5). Bound proteins were eluted by heating at 65°C for 10 min in 2× Laemmli sample buffer and analyzed by immunoblotting.

For AAL-biotin detection of protein *O*-fucosylation, protein extracts were resolved by SDS– PAGE, transferred to PVDF membranes, and probed sequentially with biotinylated AAL (1:10,000; Vector Laboratories) for 1 h and horseradish peroxidase-conjugated streptavidin for 1 h (1:5,000; Thermo Scientific).

### Transient expression in *N. benthamiana* and co-immunoprecipitation assays

*Agrobacterium tumefaciens* GV3101 strains carrying the indicated constructs were resuspended in infiltration buffer containing 10 mM MES (pH 5.6), 10 mM MgCl₂, 0.01% (v/v) Tween 20, and 150 μM acetosyringone to an OD_600_ of 1.0. The suspensions were combined as indicated for each experiment and incubated for 1 h at room temperature before infiltration into the abaxial side of approximately 6-week-old *N. benthamiana* leaves using a 1 mL needleless syringe. Leaf tissue was harvested 3 d after infiltration for protein extraction.

Co-immunoprecipitation assays were performed as described previously (Zhang et al., 2022). Approximately 200 mg of infiltrated leaf tissue was ground to a fine powder in liquid nitrogen and extracted in ice-cold lysis buffer containing 50 mM Tris-HCl (pH 7.5), 150 mM NaCl, 5 mM EDTA (pH 8.0), 0.1% (v/v) Triton X-100, and 0.2% (v/v) NP-40, supplemented immediately before use with 2 mM phenylmethylsulfonyl fluoride and 1× protease inhibitor cocktail (Roche, 11873580001). After centrifugation for 10 min at 4°C, an aliquot of each clarified extract was retained as the input, and the remaining extract was incubated with anti-GFP magnetic beads (SM038005, Smart Lifesciences) for 1.5 h at 4°C. The beads were washed three times with lysis buffer, and bound proteins were eluted by boiling in Laemmli sample buffer for 10 min. Input and immunoprecipitated fractions were analyzed by immunoblotting with anti-GFP and anti-mCherry/RFP antibodies, as indicated.

### Recombinant protein expression and purification

Recombinant 6×His–SUMO–PCK1, 6×His–SUMO–SPY, 6×His–SUMO–3TPR-SPY [The catalytic domain and three TPR domains of SPY (residues 326–914)], and GST–BIN2 were expressed and purified as described previously (Zhang et al., 2026). Briefly, the proteins were expressed in Rosetta (DE3) *Escherichia coli* cells by overnight induction with 300 μM IPTG at 16°C for 6×His–SUMO-tagged proteins or 22°C for GST-tagged proteins. Cells were lysed by ultrasonication in 1x PBS supplemented with 1 mM PMSF and 1x protease inhibitor cocktail. After clarification by centrifugation, 6×His–SUMO- and GST-tagged proteins were purified using Ni–NTA agarose (R901-01, Invitrogen) and glutathione agarose (16100, Pierce), respectively, followed by size-exclusion chromatography on a Superdex 200 column (GE Healthcare). Purified proteins were concentrated, aliquoted, and stored at −80°C until use.

### Bio-layer interferometry

The interaction between 3TPR-SPY and PCK1 was measured using a GatorPlus biolayer interferometry system (Gator Bio). 3TPR-SPY was biotinylated with EZ-Link NHS-PEG4-Biotin (Thermo A39259) and immobilized on streptavidin-XT biosensors (Gator Bio 160029) and dipped into PCK1 at 0.125, 0.25, 0.5, and 1 μM for 360 s to record association. The sensor was then dipped into PBS without PCK1 for 360 s to record dissociation. Biosensors loaded with NHS-PEG4-Biotin quenched in 50 mM Tris-Cl (pH = 7.5) was used as a control for nonspecific binding of PCK1 to the sensor surface. The dissociation constant (*K*d) was calculated using Gator Bio analysis software based on classical kinetic fitting (*k*off/*k*on).

### *In vitro O*-fucosylation assay

For *in vitro O*-fucosylation assays, 5 μg purified recombinant PCK1 protein was incubated with 1 μg recombinant SPY or 3TPR-SPY protein in 1× PBS (pH 7.3) containing 5 mM MgCl₂ and 1 mM GDP-fucose at room temperature for indicated periods or overnight, as described previously (Aizezi et al., 2024). Parallel reactions lacking SPY or GDP-fucose, or containing SPY inhibitor SOFTI, served as controls.

### Semi-*in vitro* BIN2 kinase assay

Semi-*in vitro* kinase assays were performed using the reaction conditions described previously with modifications (Zhang et al., 2026). Briefly, PCK1-YFP was immunoprecipitated with anti-GFP magnetic beads from *N. benthamiana* leaves coexpressing SPY-RFP or the RFP control. Bead-bound PCK1-YFP was incubated with 1.0 μg of purified GST-BIN2 in kinase reaction buffer containing 25 mM Tris-HCl (pH 7.4), 12 mM MgCl₂, 1 mM DTT, and 1 mM ATP at 25°C for 3 h. PCK1 phosphorylation was analyzed by Phos-tag SDS–PAGE followed by immunoblotting with anti-PCK1 antibody.

### Nuclear protein extraction

For isolation of nuclear proteins, approximately 350 mg of frozen Arabidopsis tissue powder was extracted in 1.5 mL of ice-cold buffer 1 (0.4 M sucrose, 10 mM Tris-Cl [pH 8.0], 5 mM dithiothreitol, 2 mM phenylmethylsulfonyl fluoride [PMSF], and 1× protease inhibitor cocktail). The homogenate was passed through a 70-μm cell strainer into a 50-mL conical tube and centrifuged at 800 × g for 3 min at 4 °C. The pellet was resuspended in 700 μL of the supernatant and centrifuged at 700 × g for 5 min at 4 °C. The pellet was resuspended in 1 mL of buffer 2 (0.25 M sucrose, 10 mM Tris-Cl [pH 8.0], 5 mM dithiothreitol, 10 mM MgCl₂, 1% [v/v] Triton X-100, 0.1 mM PMSF, and 1× protease inhibitor cocktail) and centrifuged at 700 × g for 5 min at 4 °C. The pellet was resuspended in 500 μL of buffer 3 (1.7 M sucrose, 10 mM Tris-Cl [pH 8.0], 5 mM dithiothreitol, 2 mM MgCl₂, 0.15% [v/v] Triton X-100, 0.1 mM PMSF, and 1× protease inhibitor cocktail), layered over a further 500 μL of buffer 3, and centrifuged at 15,000 × g for 10 min at 4 °C. The nuclear pellet was resuspended in 60 μL of 2× SDS sample buffer and heated at 65 °C for 10 min before immunoblot analysis.

### Mass spectrometric analysis of PCK1 *O*-fucosylation

PCK1-YFP was immunoprecipitated from *N. benthamiana* leaves coexpressing SPY-RFP, resolved by SDS–PAGE, and visualized by Coomassie Brilliant Blue staining. Bands corresponding to PCK1-YFP were excised, reduced, alkylated, and digested in-gel with trypsin. Peptides were analyzed using an Orbitrap Eclipse Tribrid mass spectrometer coupled to an Easy-nLC 1200 system, as described previously (Zhang et al., 2026). MS/MS spectra were searched using Protein Prospector (v6.4.9) against the *Arabidopsis thaliana* TAIR10 database concatenated with randomized decoy sequences. Carbamidomethylation of cysteine was specified as a fixed modification, whereas methionine oxidation, protein N-terminal acetylation, peptide N-terminal glutamine conversion to pyroglutamate, *O*-fucosylation of serine/threonine, and neutral loss of *O*-fucose were included as variable modifications. Precursor and fragment ion mass tolerances were set to 10 and 20 ppm, respectively, and the false discovery rate was controlled at 1% for both peptides and proteins. Up to two missed cleavages and three variable modifications per peptide were allowed.

### Statistical analysis

Statistical analyses were performed as described previously (Zhang and Kramer, 2018) using IBM SPSS Statistics (version 29.0.2.0). Data are presented as mean ± s.d. or as box plots, as indicated in the figure legends. Comparisons between two groups were performed using two-tailed Student’s *t*-test. Comparisons among multiple groups were performed using one-way ANOVA followed by Tukey’s or Games–Howell multiple-comparison test. Different letters or asterisks indicate significant differences according to the *P* values specified in the figure legends. Sample sizes and numbers of biological replicates are provided in the legends.

### Accession numbers

Sequence data from this article can be found in the GenBank/EMBL data libraries under the following Arabidopsis thaliana locus identifiers: *PCK1* (AT4G37870), *SPY* (AT3G11540), *SEC* (AT3G04240), *BIN2* (AT4G18710), *BRI1* (AT4G39400), *BZR1* (AT1G75080), and *BES1* (AT1G19350).

## Results

### SPY associates with and *O*-fucosylates PCK1

To investigate whether PCK1 is modified by SPY, we first analyzed the *O*-fucosylation of PCK1 in Arabidopsis. We crossed the 35S:PCK1-YFP line with *spy-4* to generate a homozygous 35S:PCK1-YFP/*spy-4* line. PCK1-YFP was enriched by *Aleuria aurantia* lectin (AAL)-agarose from 35S:PCK1-YFP/Col-0 extracts but not from the *spy-4* background, despite similar PCK1-YFP abundance in the inputs (Fig. 1a), indicating SPY-dependent *O*-fucosylation of PCK1 *in vivo*. Mass spectrometry detected *O*-fucosylation within an N-terminal PCK1 peptide containing Ser-62, Thr-65, and Thr-66 (Fig. 1b). Ser-62 and Thr-66 are known BIN2 phosphorylation sites (Zhang et al., 2026). Co-immunoprecipitation from *Nicotiana benthamiana* leaves showed that PCK1-YFP associates with SPY-RFP but not the RFP control (Fig. 1c). The S62A/T66A substitutions did not abolish this association.

**Figure 1:**
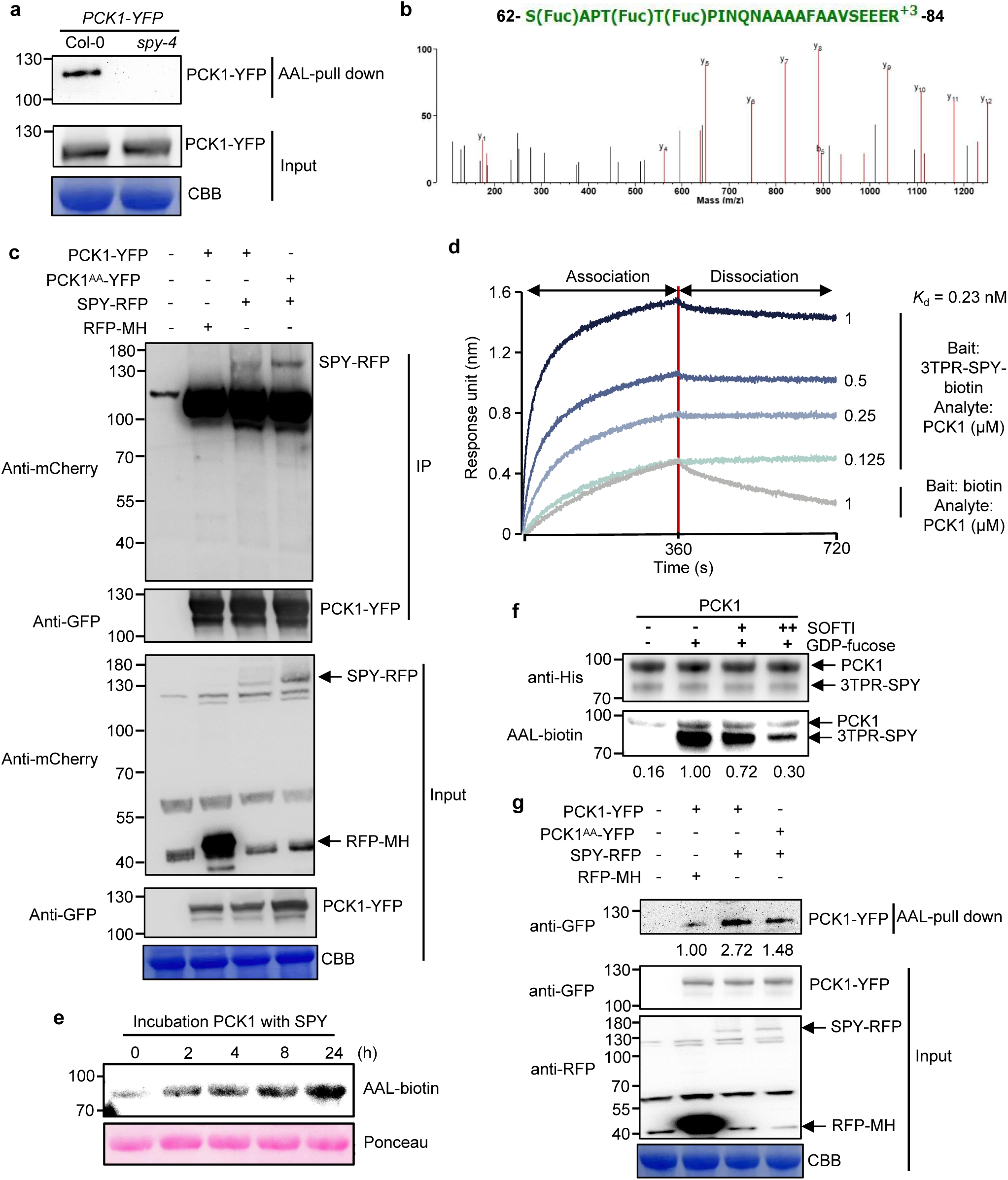
SPY associates with and *O*-fucosylates PCK1. **a**, Detection of *O*-fucosylated PCK1-YFP in PCK1-YFP/Col-0 and PCK1-YFP/*spy-4* seedlings. *O*-fucosylated proteins were pulled down by AAL-agarose. Immunoblotting of input (total protein extracts) or AAL-agarose-enriched samples was probed with an anti-GFP antibody. **b,** MS/MS spectra of fucosylated peptide 62-SAPTTPINQNAAAAFAAVSEEER-84 of PCK1-YFP protein immunoprecipitated from *N. benthamiana* leaves co-expressing PCK1-YFP with SPY-RFP, containing Ser-62, Thr-65, and Thr-66 residues. **c,** PCK1-YFP was co-immunoprecipitated with SPY-RFP in *N. benthamiana* leaves. PCK1-YFP or PCK1^AA^-YFP was transiently expressed with SPY-RFP or RFP-MH in *N. benthamiana* leaf. Immunoprecipitation was performed with total protein extracts (input) using anti-GFP beads. Input and immunoprecipitates (IPs) were analyzed by anti-mCherry and anti-GFP antibodies. **d,** Biolayer interferometry (BLI) analysis of the interaction between 3TPR-SPY and PCK1. Biotinylated 3TPR-SPY was immobilized on streptavidin biosensors and exposed to the indicated concentrations of PCK1 for 360 s, followed by dissociation in PBS for 360 s. Biosensors loaded with free biotin and exposed to 1 μM PCK1 served as a negative control. The (*K*_d_) was 0.23 nM. **e,** Time-course analysis of SPY-mediated *O*-fucosylation of recombinant PCK1. **f,** In vitro *O*-fucosylation of PCK1 by the SPY catalytic region (3TPR– SPY) by using 1 mM GDP-fucose with 10 μM (+) or 30 μM (++) of SOFTI. Fucosylation level of PCK1 and 3TPR-SPY was detected by immunoblotting with a biotinylated AAL. PCK1 and 3TPR-SPY were detected by immunoblotting with an anti-His antibody. **g,** PCK1-YFP was *O*-fucosylated by SPY in *N. benthamiana* leaves. PCK1-YFP or PCK1^AA^-YFP was transiently co-expressed with SPY-RFP or RFP-MH in *N. benthamiana* leaves. *O*-fucosylated proteins were enriched by AAL-agarose. Immunoblot containing input or AAL-agarose-enriched samples was probed with an anti-GFP or anti-RFP antibody. Values below the blots represent relative band intensities. Coomassie Brilliant Blue (CBB) or Ponceau S staining of the gel blot was used as a loading control. **Alt text:** Panel a shows lectin-enriched PCK1 immunoblots, with a band in wild type but not *spy-4*. Panel b is a fucosylated peptide mass spectrum plotting fragment intensity against mass-to-charge ratio. Panel c shows SPY co-precipitation blots for PCK1 and PCK1^AA^. Panel d plots binding response against time, with larger responses at higher PCK1 concentrations. Panel e shows progressively stronger PCK1 fucosylation bands over time. Panel f shows fucosylation blots with weaker bands at higher SOFTI concentrations. Panel g shows lectin-enriched PCK1 blots, with weaker bands for the PCK1^AA^ under SPY coexpression.

Biolayer interferometry (BLI) using purified PCK1 and a truncated SPY protein containing three tetratricopeptide repeat motifs and the catalytic region (3TPR-SPY) further supported direct binding, with an apparent dissociation constant (*K*d) of 0.23 nM (Fig. 1d). Incubation of recombinant PCK1 with SPY and GDP-fucose produced a time-dependent increase in AAL-detectable *O*-fucosylation (Fig. 1e), whereas the SPY inhibitor SOFTI reduced *O*-fucosylation of both PCK1 and SPY (Fig. 1f). In *N. benthamiana*, SPY-RFP increased AAL enrichment of PCK1-YFP more strongly than of PCK1^AA^-YFP (S62A/T66A), although the mutant retained detectable enrichment (Fig. 1g). Thus, SPY directly associates with and *O*-fucosylates PCK1, and Ser-62/Thr-66 substitutions reduce, but do not eliminate SPY-dependent PCK1 *O*-fucosylation.

### SPY-mediated *O*-fucosylation attenuates BIN2-dependent PCK1 phosphorylation

The proximity of the *O*-fucosylated PCK1 region to the known BIN2 phosphorylation sites raised the possibility that SPY-dependent *O*-fucosylation interferes with PCK1 phosphorylation. To examine this relationship, we transiently co-expressed PCK1-YFP, BIN2-YFP, and SPY-YFP in *N. benthamiana* leaves. BIN2-YFP increased PCK1-YFP phosphorylation, whereas co-expression of SPY-YFP reduced the phosphorylated fraction without appreciably changing total PCK1-YFP abundance (Fig. 2a). We then immunopurified PCK1-YFP from leaves co-expressing either SPY-RFP or an RFP control and assayed its subsequent phosphorylation by recombinant GST-BIN2 *in vitro*. PCK1-YFP isolated from SPY-expressing tissue showed substantially less BIN2-dependent phosphorylation than PCK1-YFP from the RFP control (Fig. 2b). These results indicate that SPY expression, associated with increased PCK1 *O*-fucosylation, attenuates subsequent BIN2-mediated phosphorylation.

**Figure 2:**
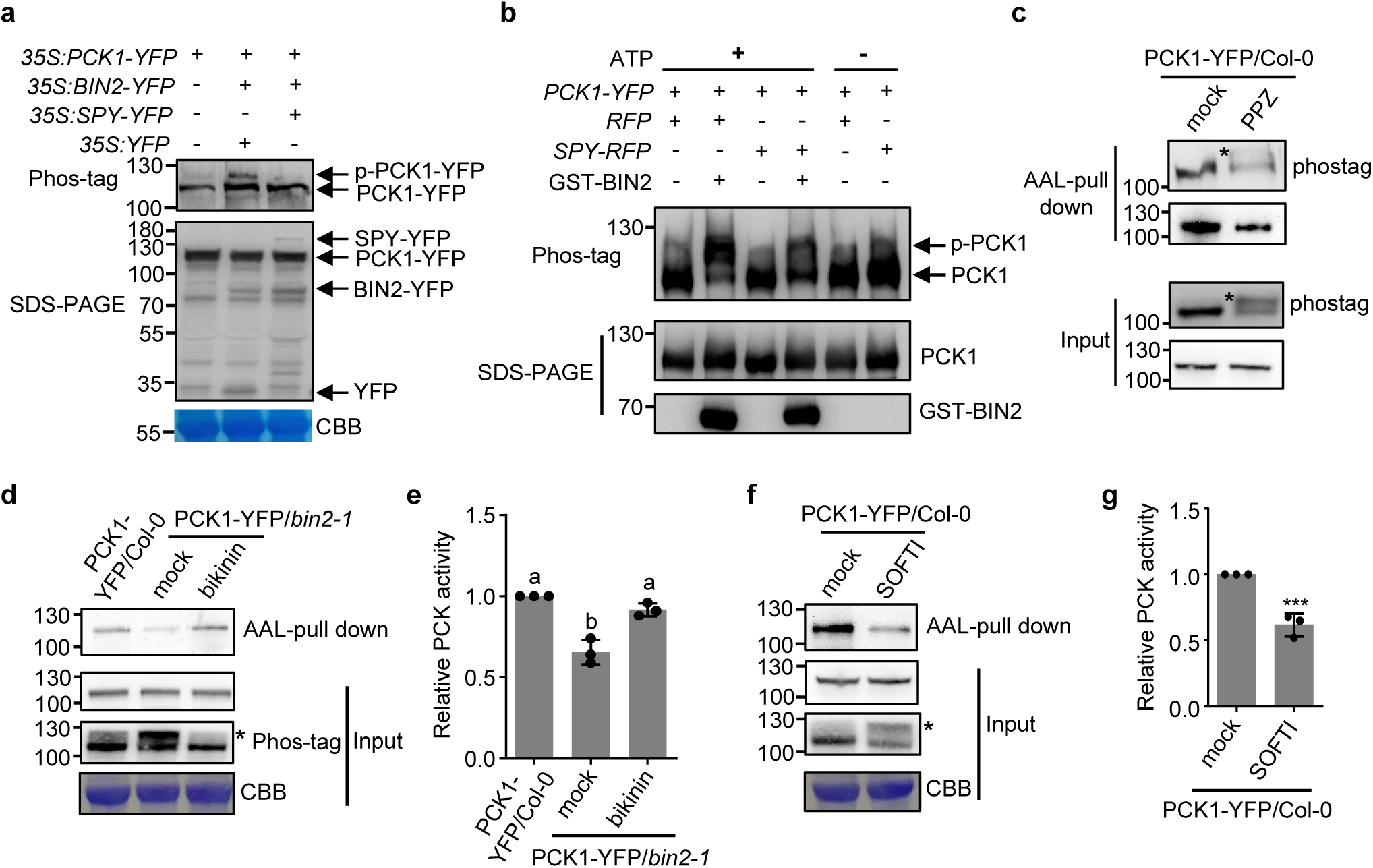
Phosphorylation and *O*-fucosylation of PCK1 antagonize each other. **a**, Co-expression of SPY with BIN2 inhibits BIN2-mediated PCK1 phosphorylation *in vivo*. The proteins were transiently co-expressed in *N. benthamiana* leaves and analyzed by immunoblotting with an anti-GFP antibody to detect protein expression by Phos-tag gel (upper blot) and standard SDS-PAGE (lower blot). **b,** PCK1 immunoprecipitated from *N. benthamiana* leaves co-expressing SPY shows reduced subsequent phosphorylation by recombinant BIN2. PCK1 was immunoprecipitated from leaves co-expressing PCK1 with SPY-RFP or RFP. We incubated the immunoprecipitated PCK1 on beads with GST-BIN2 in kinase buffer, then immunoblotted with an anti-PCK1 antibody. **c,** Immunoblotting detection of total, phosphorylated, and *O*-fucosylated PCK1 in the 7-day-old *PCK1-YFP/Col-0* transgenic seedlings grown on 0.5x MS sugar-free medium with or without 2 μM PPZ. Anti-GFP immunoblotting with input (total protein extract) or AAL-agarose pulled down samples was performed to analyze PCK1 phosphorylation status using SDS-PAGE gel containing Phos-tag (upper panel) or no Phos-tag (lower). **d,** Immunoblotting detection of total, phosphorylated, and *O*-fucosylated PCK1 in the 16-day-old seedlings of *PCK1-YFP/Col-0* and *PCK1-YFP/bin2-1* grown on 0.5x MS with 1% sucrose with or without 30 μM bikinin. **e,** Relative PCK activity of seedlings in (**d**). **f,** Immunoblotting detection of total, phosphorylated, and *O*-fucosylated PCK1 in the 16-day-old seedlings of *PCK1-YFP/Col-0* on 0.5x MS with 1% sucrose with or without 30 μM SOFTI. Anti-GFP immunoblotting with AAL-agarose-enriched samples to analyze PCK1 fucosylation or input (total protein extract) was performed to analyze PCK1 phosphorylation status using SDS-PAGE gel containing Phos-tag (lower blot) or no Phos-tag (upper blot). Phosphorylated PCK1 is the shifted band marked with *. CBB: Coomassie Brilliant Blue staining of the gel blot. **g,** Relative PCK activity of seedlings in (**f**). Data are mean ± s.d., *n* = 3 (**e,g**); ***, *P* < 0.001, according to two-tailed Student’s *t*-test (**g**), (**e**) Significant differences between means are marked by different letters (one-way ANOVA, Tukey’s test, *P* < 0.05). **Alt text:** Panel a shows PCK1 immunoblots with different combinations of BIN2 and SPY. Panel b shows kinase-assay blots with weaker phosphorylated bands after SPY coexpression. Panel c shows stronger phosphorylated and weaker lectin-enriched PCK1 bands with propiconazole. Panel d shows weaker phosphorylated and stronger lectin-enriched bands with bikinin in *bin2-1*. Panel e shows higher relative PCK activity with bikinin in a bar graph. Panel f shows stronger phosphorylated and weaker lectin-enriched bands with the SPY inhibitor SOFTI. Panel g shows lower activity with SOFTI in a bar graph.

We next examined how BR signaling and SPY activity influence PCK1 modifications and enzymatic activity in *Arabidopsis*. Inhibition of BR biosynthesis with PPZ increased PCK1 phosphorylation and decreased the amount of PCK1-YFP recovered by AAL-agarose without markedly changing total PCK1-YFP abundance (Fig. 2c). The phosphorylated PCK1 fraction was lower in the AAL-enriched material than in the corresponding input, consistent with an inverse relationship between the two modifications (Fig. 2c). In PCK1-YFP/*bin2-1* seedlings, where BIN2 activity is elevated, PCK1 phosphorylation increased while AAL enrichment decreased relative to the WT background (Fig. 2d). Bikinin, a BIN2/GSK3 inhibitor, reduced PCK1 phosphorylation, increased AAL enrichment, and partly restored PCK activity, which was approximately 65% of WT in *bin2-1* without treatment (Fig. 2d,e). We also examined SPY inhibition in PCK1-YFP/Col-0 seedlings. SOFTI decreased AAL enrichment of PCK1-YFP without a marked change in its total abundance, accompanied by greater PCK1 phosphorylation and a reduction of PCK activity to approximately 60% of the control level (Fig. 2f,g). Together, these experiments support reciprocal effects of SPY-dependent *O*-fucosylation and BIN2-dependent phosphorylation on PCK1 modification status and activity.

### Loss of SPY reduces PCK1 activity by increasing PCK1 phosphorylation

We next asked whether increased phosphorylation contributes to the reduced PCK activity observed in *spy* seedlings. In 5-d-old seedlings, *spy-4* and *spy-23* each showed approximately half the PCK activity of WT, together with a higher proportion of phosphorylated PCK1 (approximately 57% versus 22% in WT; Fig. 3a). Treatment with 1 nM BL lowered PCK1 phosphorylation to approximately 17–18% and restored PCK activity in both *spy* alleles toward the WT level (Fig. 3a). In 3.5-d-old etiolated seedlings grown on sugar-free medium, basal PCK1 phosphorylation was approximately 24–26% in WT and 44– 47% in *spy* mutants, accompanied by lower PCK activity in the mutants (Fig. 3b,c). PPZ increased PCK1 phosphorylation to approximately 51–54% in WT and 67–69% in *spy* mutants and further lowered PCK activity. Addition of either BL or bikinin reduced the PPZ-associated phosphorylated fraction and restored activity substantially in both genotypes (Fig. 3b,c). These results indicate that the PCK activity defect in *spy* seedlings is reversible by inhibiting BIN2-dependent phosphorylation through BR signaling or a kinase inhibitor.

**Figure 3:**
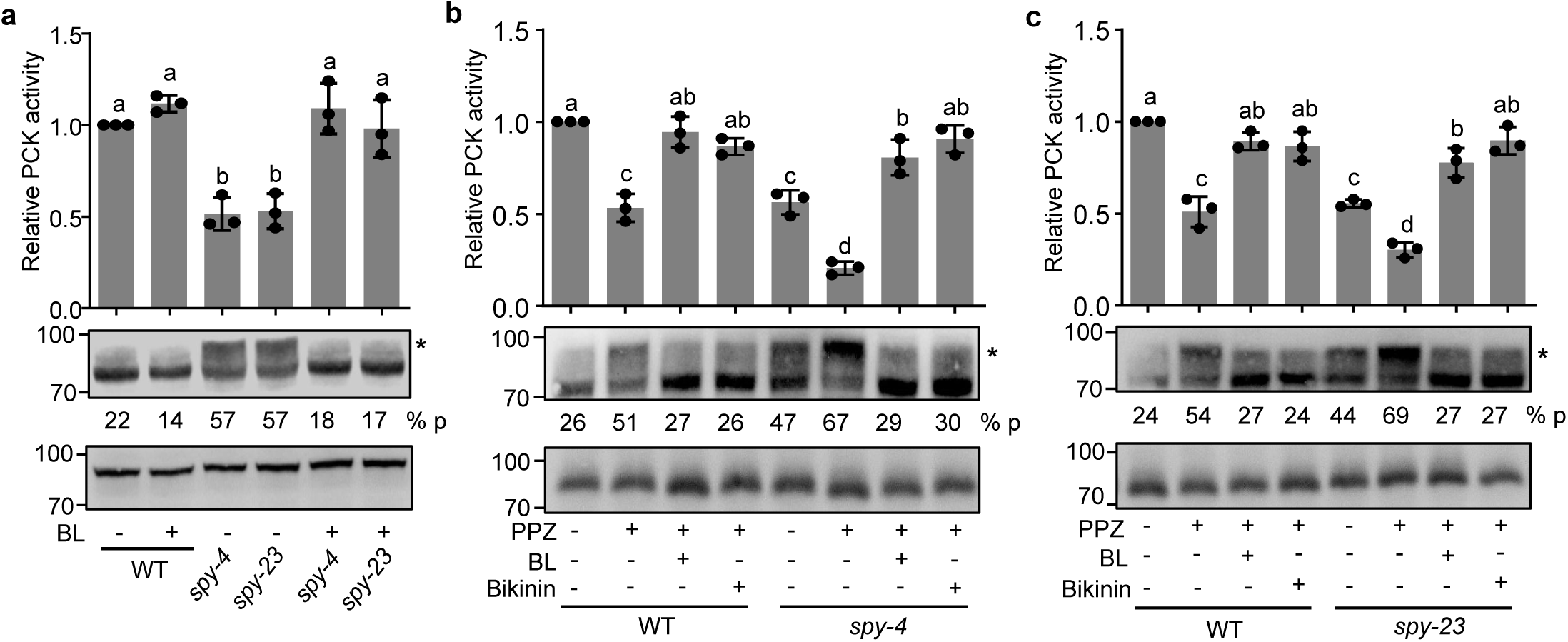
The *spy* mutants show increased PCK1 phosphorylation and reduced PCK activity. **a**, Relative PCK activity and phosphorylation status of WT and *spy* mutants grown on medium with or without 1 nM brassinolide (BL) for 5 days. **b-c,** Relative PCK decarboxylation activity and PCK1 phosphorylation status of 3.5-day-old WT and *spy* etiolated seedlings grown on 0.5x MS sugar-free medium containing 2 μM propiconazole (PPZ), 2 μM PPZ plus 100 nM BL, or 2 μM PPZ plus 30 μM bikinin. PCK1 phosphorylation status was analyzed using anti-PCK1 immunoblotting of SDS-PAGE gel containing Phos-tag (upper panel) or no Phos-tag (lower panel). Phosphorylated PCK1 is the shifted band marked with *. Values below the blots represent the percentage of phosphorylated PCK1 (% p) relative to total PCK1. Data are mean ± s.d., *n* = 3 biological replicates (**a-c**); Significant differences between means are marked by different letters (one-way ANOVA, Tukey’s test, *P* < 0.05). **Alt text:** Panel a shows relative PCK activity bars above PCK1 immunoblots for wild type and two *spy* mutants with or without brassinolide. Untreated mutants have lower activity and stronger phosphorylated bands. Brassinolide raises activity and weakens these bands. Panel b compares wild type and *spy-4* under propiconazole alone or with brassinolide or bikinin; combined treatments raise activity and weaken phosphorylated bands. Panel c shows activity bars and blots for wild type and *spy-23*; combined treatments likewise raise activity and weaken phosphorylated bands.

### *O*-fucosylation promotes seedling growth and is inhibited by BIN2

To test whether loss of *O*-fucosylation enhances BIN2-dependent phosphorylation and thus contributes to growth inhibition, we analyzed the phenotypes of *spy* mutants in the dark. On sugar-free medium, *pck1* and *spy* showed short hypocotyls (Fig. 4a,b). On media containing exogenous sucrose, the *pck1* phenotype was rescued (Fig. 4a,b), as reported previously. However, sucrose did not rescue the short-hypocotyl phenotype of *spy-4* or *spy*-23 (Fig. 4a,b). Dysfunction in additional SPY substrates likely contributes to the growth defects, as PCK1 is one of hundreds of proteins targeted by SPY.

**Figure 4:**
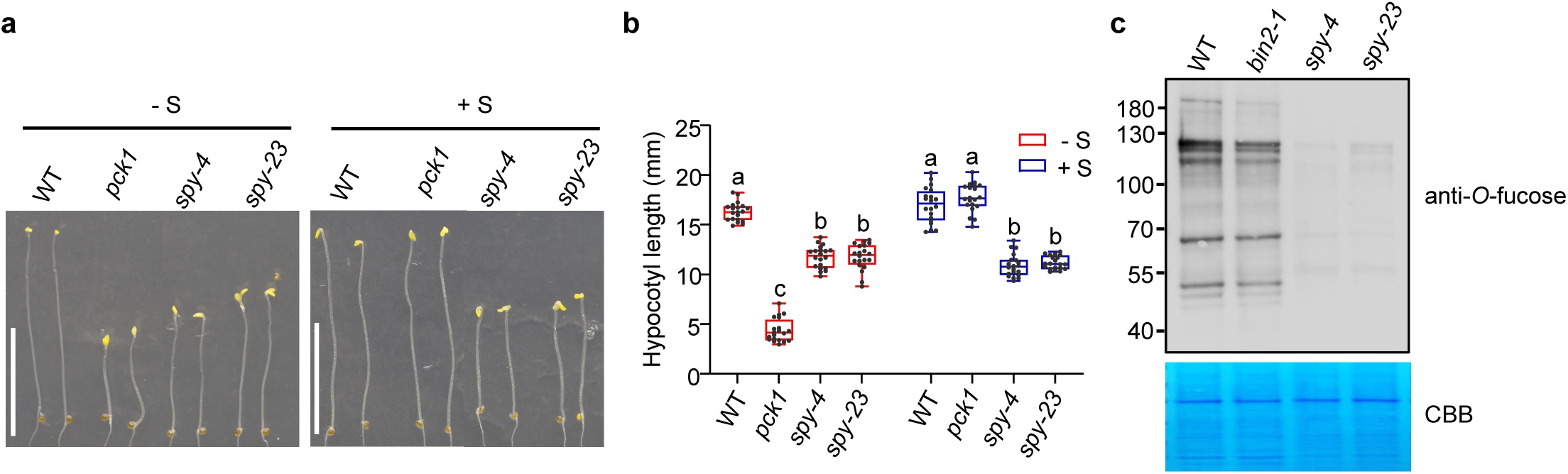
*O*-fucosylation promotes seedling growth and is inhibited by BIN2. **a**, Photograph of 5-day-old WT, *pck1*, *spy-4*, and *spy-23* seedlings grown on 0.5x MS medium with or without 1% sucrose. Scale bars, 10 mm. **b,** Hypocotyl length of seedlings in (**a**). *n* = 20 seedlings. The central lines within the box plot represent the medians, the box represents the interquartile range (IQR), and the whiskers extend to the minima and maxima. Significant differences between means are marked by different letters (one-way ANOVA, Games– Howell test, *P* < 0.05). **c,** Detection of *O*-fucosylated proteins in WT, *bin2-1*, and *spy* mutants by immunoblotting. Total nuclear proteins were extracted and immunoblotted with a mixture of anti-Serine-*O-*fucose and anti-Threonine-*O-*fucose antibodies. CBB: Coomassie Brilliant Blue staining of the gel blot. **Alt text:** Panel a shows photographs of wild-type, *pck1*, *spy-4* and *spy-23* seedlings with or without sucrose. Panel b shows box plots and individual data points for hypocotyl length; sucrose-treated *pck1* seedlings have lengths similar to wild type, while both *spy* mutants remain shorter. Panel c shows an immunoblot of *O*-fucosylated nuclear proteins; several *bin2-1* bands are fainter than wild type, and *spy* bands are barely visible.

The significant overlap between targets of SPY and BIN2 raises the possibility that SPY antagonizes BIN2 and enhances BR-regulated processes globally. To explore whether BIN2 might also influence *O*-fucosylation of proteins other than PCK1, we examined nuclear protein fractions with antibodies recognizing serine- and threonine-linked *O*-fucose. Several bands showed lower intensity in the gain-of-function mutant *bin2-1* than in WT, and were not detected in the *spy* mutants (Fig. 4c). These observations indicate that BIN2 reduces *O*-fucosylation of multiple proteins in addition to PCK1.

### SPY promotes BR response in dark-grown seedlings

To test this possibility, we analyzed the BR responses of *spy* mutants. The *spy* mutants displayed shorter hypocotyls than wild type under normal conditions but showed no significant difference from wild type when grown on 2 µM PPZ, or PPZ plus 100 nM brassinolide (BL) or 30 µM bikinin (Fig. 5a,b), indicating that the short-hypocotyl phenotype of *spy* mutants is dependent on BR. We then grew the mutants on various concentrations of PPZ and 2 µM PPZ plus various concentrations of BL. The results show that the *spy-4* and *spy-23* mutants were hypersensitive to PPZ treatments, in contrast to *bzr1-1D,* which is hyposensitive (Fig. 5c,d), but were hyposensitive to exogenous BL: WT reached maximal elongation on 10 nM BL, whereas both *spy* alleles continued to increase hypocotyl length as BL concentration increased to 100 nM (Fig. 5e). These results suggest that sugar-dependent *O*-glycosylation promotes plant growth in part by antagonizing BIN2-dependent phosphorylation.

**Figure 5:**
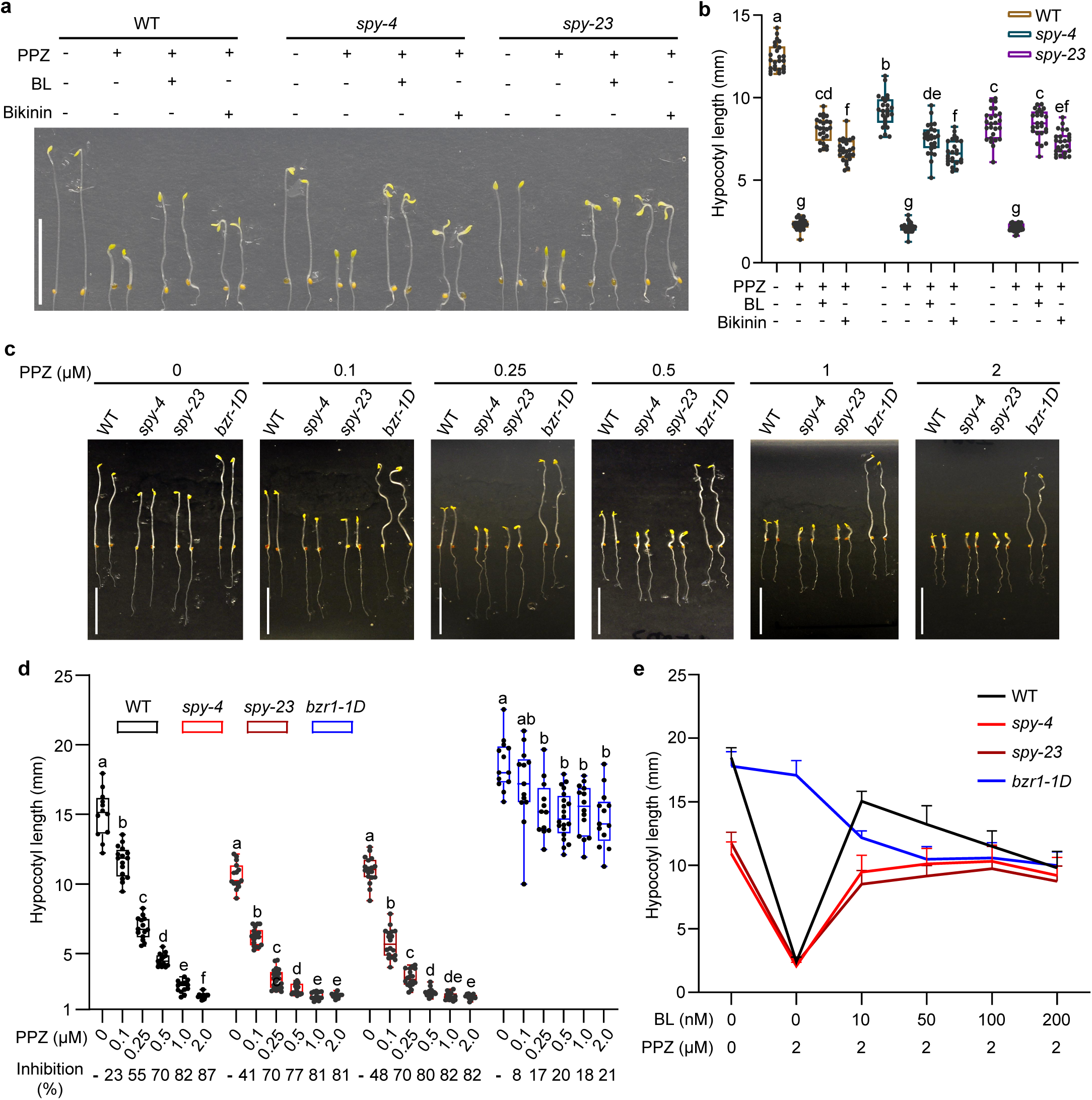
SPY promotes BR response in dark-grown seedlings. **a**, Photograph of 3.5-day-old WT and *spy* seedlings grown in the dark on 0.5x MS sugar-free medium containing 2 μM propiconazole (PPZ), 2 μM PPZ plus 100 nM brassinolide (BL), or 2 μM PPZ plus 30 μM bikinin. Scale bar, 10 mm. **b,** Hypocotyl length of seedlings shown in (**a**). *n* = 25 seedlings. **c,** Photograph of 5-day-old WT, *spy* mutants, and *bzr1-1D* seedlings grown in the dark on 0.5x MS medium with 1% sucrose containing different concentrations of PPZ in the dark. Scale bars, 10 mm. **d,** Hypocotyl length of seedlings in (**c**). *n* ≥ 12 seedlings. The central lines within the box plots (**b,d**) represent the medians, the box represents the interquartile range (IQR), and the whiskers extend to the minima and maxima. Percentage values indicate the relative reduction in hypocotyl length at each PPZ concentration compared with the untreated control of the same genotype. Significant differences between means (**b**) or between means within each genotype (**d**) are marked by different letters (one- way ANOVA, Tukey’s test, **b**; Games–Howell test, **d**; *P* < 0.05). **e,** Hypocotyl length of 5-day-old seedlings of WT, *spy*, and *bzr1-1D* mutants grown on 0.5x MS medium or 0.5x MS medium containing 2 μM PPZ plus various concentrations of BL. Data are mean ± s.d., *n* = 12. **Alt text:** Panel a shows photographs of wild-type and *spy* seedlings. Panel b shows box plots of hypocotyl length with individual points; adding brassinolide or bikinin to propiconazole increases length. Panel c shows photographs of wild-type, *spy* and *bzr1-1D* seedlings across propiconazole doses. Panel d shows corresponding box plots; low doses shorten *spy* proportionally more than wild type, while *bzr1-1D* remains longest. Panel e plots hypocotyl length across brassinolide concentrations, including untreated controls. Under propiconazole, increasing brassinolide produces a rise then fall in wild type, a smaller rise in *spy* and a decline in *bzr1-1D*; lengths converge at higher doses.

## Discussion

Plant growth requires coordination between carbon availability and hormone signaling. Nucleocytoplasmic *O*-glycosylation links nutrient status to protein regulation, while BIN2-dependent phosphorylation connects brassinosteroid (BR) signaling to numerous cellular targets (Bi et al., 2023; Kim et al., 2023; Aizezi et al., 2025). To investigate how these modifications interact to regulate protein function, we examined SPY-mediated *O*-fucosylation of PCK1, a key enzyme in carbon metabolism whose activity is directly inhibited by BIN2 phosphorylation (Zhang et al., 2026). Our findings demonstrate reciprocal antagonism between PCK1 *O*-fucosylation and phosphorylation and show that SPY promotes PCK1 activity by counteracting BIN2-dependent inhibition. PCK1 thus provides a mechanistic connection between sugar-responsive *O*-fucosylation and BR-regulated phosphorylation in the control of carbon metabolism.

The reduced PCK1 activity in the *spy* mutant supports a positive role for SPY-mediated *O*-fucosylation in PCK1 regulation in vivo. Mass spectrometry detected *O*-fucosylation within an N-terminal peptide containing Ser-62, Thr-65, and Thr-66. Ser-62 and Thr-66 are established BIN2 phosphorylation sites, and our previous work showed that phosphorylation at these residues inhibits PCK1 by shifting the enzyme from an active hexameric form to an inactive dodecameric form (Zhang et al., 2026). Substitution of Ser-62 and Thr-66 with alanine reduced SPY-dependent *O*-fucosylation, consistent with overlapping sites of *O*-fucosylation and phosphorylation within this regulatory region. These observations raise two possible mechanisms by which *O*-fucosylation could promote PCK1 activity: directly influencing enzyme structure or preventing the inhibitory phosphorylation that alters its oligomeric state.

Genetic and pharmacological perturbations further support reciprocal antagonism between the two modifications *in vivo*. SPY overexpression reduced BIN2-mediated PCK1 phosphorylation, whereas loss of SPY or its chemical inhibition by SOFTI caused PCK1 hyperphosphorylation. Conversely, promoting BIN2 activity by inhibiting BR biosynthesis decreased PCK1 *O*-fucosylation, whereas bikinin treatment increased it. AAL lectin preferentially enriched unphosphorylated PCK1, further supporting an inverse relationship between *O*-fucosylation and phosphorylation. Together with the mass spectrometry and mutational data, these findings support a model in which the two modifications compete at shared or closely spaced residues within the N-terminal regulatory region. Functional rescue of *spy* provides further insight into the consequences of this interplay. Treatment with BL or bikinin reversed PCK1 hyperphosphorylation and restored its activity to wild-type levels, demonstrating that relief of inhibitory phosphorylation is sufficient to recover PCK1 activity in the absence of SPY. These results support protection from BIN2-dependent inhibition as a principal mechanism through which SPY-mediated *O*-fucosylation promotes PCK1 activity.

The SPY–BIN2 antagonism demonstrated for PCK1 may extend to other cellular proteins, suggesting a broader regulatory relationship between *O*-glycosylation and GSK3-mediated phosphorylation. Dark-grown *spy* seedlings exhibit short hypocotyls, resembling *pck1* mutants and consistent with reduced PCK1 activity. However, exogenous sugar rescues the short-hypocotyl phenotype of *pck1* but not that of *spy*, suggesting that additional SPY targets contribute to the growth defects of *spy*. Previous proteomic studies identified more than 100 BIN2 substrates that are also modified by *O*-GlcNAc and/or *O*-fucose, highlighting extensive potential for crosstalk between *O*-glycosylation and phosphorylation. Consistent with this possibility, our anti-*O*-fucose immunoblot analysis revealed an overall reduction in protein *O*-fucosylation in the dominant *bin2-1* mutant, suggesting that BIN2-dependent regulation of *O*-fucosylation extends beyond PCK1. Broader antagonism between SPY and BIN2 may therefore contribute to the short-hypocotyl and BR-hyposensitive phenotypes of dark-grown *spy* seedlings. Suppression of these phenotypes by high concentrations of BL or by bikinin further supports involvement of the BR–BIN2 pathway in these growth defects. Together, our findings establish reciprocal antagonism between *O*-fucosylation and phosphorylation as a mechanism regulating PCK1 and suggest that similar interactions among shared substrates help coordinate carbon metabolism and plant growth. Functional characterization of additional SPY–BIN2 substrates will clarify the scope and physiological significance of this mechanism in integrating nutrient and hormone signaling.

## Author contributions

H.Z. and Z.W. designed the research. H.Z., Y.A., and A.C. performed experiments. H.Z. and

Z.W. analyzed data. H.Z. and Z.W. wrote and edited the manuscript.

## Acknowledgement

The authors thank Alberto A. Iglesias (Instituto de Agrobiotecnología del Litoral, UNL, CONICET, FBCB, Santa Fe, Argentina) for sharing the anti-PCK1 antibody, and Drs. Cao Son Trinh and Shouling Xu for assistance with mass spectrometry analyses.

## Data availability statement

All materials and data presented in this study are available from the corresponding author on request.

## References

Aizezi, Y., Yuan, Y., Xu, S.L., and Wang, Z.Y. (2025). A tale of two sugars: O-GlcNAc and O-fucose orchestrate growth, development, and acclimation in plants. Trends Biochem Sci 50, 332–343.

Aizezi, Y., Zhao, H., Zhang, Z., Bi, Y., Yang, Q., Guo, G., Zhang, H., Guo, H., Jiang, K., and Wang, Z.Y. (2024). Structure-based virtual screening identifies small-molecule inhibitors of O-fucosyltransferase SPINDLY in Arabidopsis. Plant Cell 36, 497–509.

Bi, Y., Deng, Z., Ni, W., Shrestha, R., Savage, D., Hartwig, T., Patil, S., Hong, S.H., Zhang, Z., Oses-Prieto, J.A., Li, K.H., Quail, P.H., Burlingame, A.L., Xu, S.L., and Wang, Z.Y. (2021). Arabidopsis ACINUS is O-glycosylated and regulates transcription and alternative splicing of regulators of reproductive transitions. Nat Commun 12, 945.

Bi, Y., Shrestha, R., Zhang, Z., Hsu, C.C., Reyes, A.V., Karunadasa, S., Baker, P.R., Maynard, J.C., Liu, Y., Hakimi, A., Lopez-Ferrer, D., Hassan, T., Chalkley, R.J., Xu, S.L., and Wang, Z.Y. (2023). SPINDLY mediates O-fucosylation of hundreds of proteins and sugar-dependent growth in Arabidopsis. Plant Cell 35, 1318–1333.

Burnell, J.N., and Hatch, M.D. (1988). Photosynthesis in phosphoenolpyruvate carboxykinase-type C4 plants: Pathways of C4 acid decarboxylation in bundle sheath cells of Urochloa panicoides. Archives of Biochemistry and Biophysics 260, 187–199.

Hart, G.W., Housley, M.P., and Slawson, C. (2007). Cycling of O-linked beta-N-acetylglucosamine on nucleocytoplasmic proteins. Nature 446, 1017–1022.

Hart, G.W., Slawson, C., Ramirez-Correa, G., and Lagerlof, O. (2011). Cross talk between O-GlcNAcylation and phosphorylation: roles in signaling, transcription, and chronic disease. Annual review of biochemistry 80, 825–858.

Hartweck, L.M., Scott, C.L., and Olszewski, N.E. (2002). Two O-linked N-acetylglucosamine transferase genes of Arabidopsis thaliana L. Heynh. have overlapping functions necessary for gamete and seed development. Genetics 161,3, 1279–1291.

He, J.-X., Gendron, J.M., Yang, Y., Li, J., and Wang, Z.-Y. (2002). The GSK3-like kinase BIN2 phosphorylates and destabilizes BZR1, a positive regulator of the brassinosteroid signaling pathway in Arabidopsis. Proceedings of the National Academy of Sciences 99, 10185–10190.

Kim, T.W., Guan, S., Sun, Y., Deng, Z., Tang, W., Shang, J.X., Sun, Y., Burlingame, A.L., and Wang, Z.Y. (2009). Brassinosteroid signal transduction from cell-surface receptor kinases to nuclear transcription factors. Nat Cell Biol 11, 1254–1260.

Kim, T.W., Park, C.H., Hsu, C.C., Kim, Y.W., Ko, Y.W., Zhang, Z., Zhu, J.Y., Hsiao, Y.C., Branon, T., Kaasik, K., Saldivar, E., Li, K., Pasha, A., Provart, N.J., Burlingame, A.L., Xu, S.L., Ting, A.Y., and Wang, Z.Y. (2023). Mapping the signaling network of BIN2 kinase using TurboID-mediated biotin labeling and phosphoproteomics. Plant Cell 35, 975–993.

Lagerlöf, O., Slocomb, J.E., Hong, I., Aponte, Y., Blackshaw, S., Hart, G.W., and Huganir, R.L. (2016). The nutrient sensor OGT in PVN neurons regulates feeding. Science 351, 1293–1296.

Leegood, R.C., and Walker, R.P. (2003). Regulation and roles of phosphoenolpyruvate carboxykinase in plants. Arch Biochem Biophys 414, 204–210.

Li, J., and Nam, K.H. (2002). Regulation of Brassinosteroid Signaling by a GSK3/SHAGGY-Like Kinase. Science 295, 1299–1301.

Penfield, S., Rylott, E.L., Gilday, A.D., Graham, S., Larson, T.R., and Graham, I.A. (2004). Reserve mobilization in the Arabidopsis endosperm fuels hypocotyl elongation in the dark, is independent of abscisic acid, and requires PHOSPHOENOLPYRUVATE CARBOXYKINASE1. Plant Cell 16, 2705–2718.

Rojas, B.E., Hartman, M.D., Figueroa, C.M., and Iglesias, A.A. (2021). Proteolytic cleavage of Arabidopsis thaliana phosphoenolpyruvate carboxykinase-1 modifies its allosteric regulation. J Exp Bot 72, 2514–2524.

Rylott, E.L., Gilday, A.D., and Graham, I.A. (2003). The gluconeogenic enzyme phosphoenolpyruvate carboxykinase in Arabidopsis is essential for seedling establishment. Plant Physiol 131, 1834–1842.

Shrestha, R., Karunadasa, S., Grismer, T.S., Reyes, A.V., and Xu, S.L. (2024). SECRET AGENT O-GlcNAcylates Hundreds of Proteins Involved in Diverse Cellular Processes in Arabidopsis. Mol Cell Proteomics 23, 100732.

Swain, S.M., Tseng, T.S., and Olszewski, N.E. (2001). Altered expression of SPINDLY affects gibberellin response and plant development. Plant Physiol 126, 1174–1185.

Walker, R.P., and Leegood, R.C. (1996). Phosphorylation of phosphoenolpyruvate carboxykinase in plants. Studies in plants with C4 photosynthesis and Crassulacean acid metabolism and in germinating seeds. Biochemical Journal 317, 653–658.

Walker, R.P., Chen, Z.-H., and Famiani, F. (2021). Gluconeogenesis in Plants: A Key Interface between Organic Acid/Amino Acid/Lipid and Sugar Metabolism. Molecules 26.

Woody, S.T., Austin-Phillips, S., Amasino, R.M., and Krysan, P.J. (2007). The WiscDsLox T-DNA collection: an arabidopsis community resource generated by using an improved high-throughput T-DNA sequencing pipeline. J Plant Res 120, 157–165.

Xu, S.L., Chalkley, R.J., Maynard, J.C., Wang, W., Ni, W., Jiang, X., Shin, K., Cheng, L., Savage, D., Huhmer, A.F., Burlingame, A.L., and Wang, Z.Y. (2017). Proteomic analysis reveals O-GlcNAc modification on proteins with key regulatory functions in Arabidopsis. Proc Natl Acad Sci U S A 114, E1536–E1543.

Zentella, R., Wang, Y., Zahn, E., Hu, J., Jiang, L., Shabanowitz, J., Hunt, D.F., and Sun, T.P. (2023). SPINDLY O-fucosylates nuclear and cytoplasmic proteins involved in diverse cellular processes in plants. Plant Physiol 191, 1546–1560.

Zentella, R., Sui, N., Barnhill, B., Hsieh, W.P., Hu, J., Shabanowitz, J., Boyce, M., Olszewski, N.E., Zhou, P., Hunt, D.F., and Sun, T.P. (2017). The Arabidopsis O-fucosyltransferase SPINDLY activates nuclear growth repressor DELLA. Nat Chem Biol 13, 479–485.

Zentella, R., Hu, J., Hsieh, W.P., Matsumoto, P.A., Dawdy, A., Barnhill, B., Oldenhof, H., Hartweck, L.M., Maitra, S., Thomas, S.G., Cockrell, S., Boyce, M., Shabanowitz, J., Hunt, D.F., Olszewski, N.E., and Sun, T.P. (2016). O-GlcNAcylation of master growth repressor DELLA by SECRET AGENT modulates multiple signaling pathways in Arabidopsis. Genes Dev 30, 164–176.

Zhang, H., and Kramer, U. (2018). Differential Diel Translation of Transcripts With Roles in the Transfer and Utilization of Iron-Sulfur Clusters in Arabidopsis. Front Plant Sci 9, 1641.

Zhang, H., Aizezi, Y., Bessho-Uehara, K., Chaudhary, A., Trinh, C.S., Xu, S.L., and Wang, Z.Y. (2026). Brassinosteroids promote sugar synthesis by inhibiting BIN2 phosphorylation of phosphoenolpyruvate carboxykinase. Nat Plants 12, 1741–1755.

Zhang, H., Janina, N., Utkur, K., Manivannan, T., Zhang, L., Wang, L., Grefen, C., Schaffrath, R., and Kramer, U. (2025). Diphthamide formation in Arabidopsis requires DPH1-interacting DPH2 for light and oxidative stress resistance. Plant Physiol 197.

Zhang, H., Quintana, J., Utkur, K., Adrian, L., Hawer, H., Mayer, K., Gong, X., Castanedo, L., Schulten, A., Janina, N., Peters, M., Wirtz, M., Brinkmann, U., Schaffrath, R., and Kramer, U. (2022). Translational fidelity and growth of Arabidopsis require stress-sensitive diphthamide biosynthesis. Nat Commun 13, 4009.

Zhang, Z., Zhu, J.Y., Roh, J., Marchive, C., Kim, S.K., Meyer, C., Sun, Y., Wang, W., and Wang, Z.Y. (2016). TOR Signaling Promotes Accumulation of BZR1 to Balance Growth with Carbon Availability in Arabidopsis. Curr Biol 26, 1854–1860.

